# Characterization of the trimeric TOM complex by HS-AFM single-molecule analysis

**DOI:** 10.64898/2026.07.01.735793

**Authors:** Nanako Kobayashi, Satoshi N. Omura, Kana Kuzasa, Kenichiro Imai, Shiho Kawai, Hirotatsu Imai, Romain Amyot, Kenichi Umeda, Osamu Nureki, Toshiya Endo, Noriyuki Kodera, Yuhei Araiso

## Abstract

The translocase of the outer mitochondrial membrane (TOM) complex is the main entry gate for mitochondrial proteins. Approximately 99 % of mitochondrial proteins are synthesized as precursor proteins (preproteins) in the cytosol and subsequently translocated into mitochondria through the TOM complex. The TOM complex exists in a dynamic equilibrium among multiple assembly states through spatial rearrangements of its subunits. The recent cryo-electron microscopy (cryo-EM) studies revealed near-atomic structures of the TOM core dimer, whereas previous biochemical studies indicated the TOM complex functions as a trimer in intact mitochondria. However, the relationship between the core dimer and the functional trimer remains unclear. In the present study, we analyzed the dynamics of the TOM complex using high-speed atomic force microscopy (HS-AFM) to investigate the assembly states and conformation transitions of the TOM complexes. We demonstrated that purified yeast TOM complexes predominantly adopt a trimeric organization but dynamically dissociate into dimeric and monomeric states during HS-AFM observation. The trimeric particles observed by HS-AFM exhibited spherical molecular shapes consistent with a trimeric structural model proposed from previous crosslinking analyses. In contrast, the dissociated dimeric particles closely resembled the dimensions of the TOM core-dimer structures determined by cryo-EM. Furthermore, HS-AFM analyses provided insight into the spatial arrangement of the Tom20 receptor, consistent with previous models of the trimeric TOM complex. These observations enabled characterization of the trimeric TOM complex *in vitro* and provide a foundation for future structural and functional analyses of TOM complex assembly.

**Significance statement:** The translocase of the mitochondrial outer membrane (TOM) complex is the universal entry gate for almost all mitochondrial proteins, yet its native organization has remained elusive. By combining high-speed atomic force microscopy with cryo-electron microscopy, we provide evidence that the yeast TOM complex predominantly adopts a trimeric assembly with three protein-conducting channels, rather than the dimeric architecture proposed by previous structural studies. Furthermore, single-molecule imaging directly visualized interactions with mitochondrial precursor proteins and receptor-specific antibodies, enabling the experimental identification of receptor subunits within the trimeric TOM complex. Together, these findings uncover previously unrecognized structural and dynamic features of the TOM complex and provide a framework for understanding the molecular mechanism of mitochondrial protein import.

## Introduction

Mitochondria are essential organelles that maintain cellular homeostasis and mediate diverse cellular functions. In humans, mitochondria contain approximately 1,500 different proteins, whereas yeast mitochondria contain approximately 800 proteins. However, only a small number of mitochondrial proteins are encoded by the mitochondrial genome. More than 99% of mitochondrial proteins are synthesized in the cytosol and subsequently imported into mitochondria, where they are sorted into the outer membrane, intermembrane space, inner membrane, and matrix compartments (1–4).

The translocase of the outer mitochondrial membrane (TOM) complex serves as the major entry gate for the import of mitochondrial precursor proteins (preproteins). The TOM complex consists of the β-barrel channel protein Tom40 and six receptor or auxiliary subunits, including Tom70, Tom22, Tom20, Tom7, Tom6, and Tom5, each of which contains a single transmembrane helix (5). Although Tom40 was identified as the central import channel of the TOM complex around 1990 (6–8), the high-resolution structure of the TOM complex remained unresolved for many years. Early electron microscopy studies using negatively stained particles demonstrated that purified TOM complexes from *Saccharomyces cerevisiae* and *Neurospora crassa* contained mixtures of trimeric and dimeric particles with three or two apparent protein-conducting pores, respectively (9,10). Subsequently, moderate-resolution cryo-electron microscopy (∼18 Å) suggested that the TOM complex can adopt a trimeric architecture and that the presence of Tom20 favors formation of the trimeric complex (11). Systematic crosslinking analyses further supported a trimeric TOM complex containing three copies each of Tom40 and Tom22 (12). Biochemical analyses also showed that the trimeric TOM complex dynamically exchanges with a minor TOM species lacking Tom22, which likely consists of dimeric Tom40 pores and mediates the import of the intermembrane space-targeted proteins (13).

More recently, near-atomic-resolution structures of the yeast TOM complex were determined by single-particle cryo-electron microscopy (14–16). Unexpectedly, these structures revealed a core dimer composed of two Tom40 channels surrounded by Tom22, Tom7, Tom6, and Tom5, which did not fully reconcile with the trimeric and Tom22-lacking dimeric forms suggested by previous biochemical studies (12,13). Soon afterward, cryo-EM structures of the human TOM complex demonstrated a similar overall architecture (17,18). Structural analyses of holo-TOM complexes have also progressively clarified the positions of Tom20 subunits that were previously unresolved; however, these structures likewise adopted dimeric organizations (19–22).

These observations raise an important question regarding the relationship between the trimeric TOM complexes observed in intact mitochondria and the core-dimer structures resolved by cryo-EM analysis. Earlier electron microscopy studies suggested that purified TOM complexes contain mixtures of trimeric and dimeric forms (9–11). In addition, previous high-speed atomic force microscopy (HS-AFM) studies suggested that purified TOM complexes can adopt trimeric structures that tended to dissociate into dimeric and monomeric forms (15). However, the detailed molecular architecture and dynamic behavior of these particles have not been fully characterized, and the relationship between the TOM complexes observed by cryo-EM and HS-AFM remains unclear.

In this study, we demonstrated that purified yeast TOM complexes predominantly adopt a trimeric organization but dynamically dissociate into dimeric and monomeric states during HS-AFM observation. The trimeric particles observed by HS-AFM exhibited spherical molecular shapes consistent with a trimeric structural model proposed from previous crosslinking analyses (12). In contrast, the dissociated dimeric particles closely matched the dimensions of the TOM core-dimer structures previously determined by cryo-EM (15). We also captured interactions between a preprotein and the TOM complex upon addition of a model substrate. Furthermore, antibody-labeling experiments targeting Tom20 provided direct evidence supporting a trimeric organization of the TOM complex.

## Results

### The HS-AFM imaging of the trimeric TOM complex

To advance the structural analysis of the TOM complex, we constructed a budding yeast strain overexpressing each TOM complex subunit, including Tom40-Strep-tag II, Tom22-10×His, Tom20, Tom7, Tom6, and Tom5. Both the Strep-tag II and 10×His tag were positioned on the intermembrane space (IMS) side of the complex. We first isolated mitochondria and affinity-purified the TOM complex using the Strep-tag II under detergent micelle conditions. Gel filtration chromatography demonstrated that the purified TOM complex eluted as a sharp peak at 14–15 mL (Fig. 1A). Immunoblot analyses following SDS-PAGE and blue-native PAGE (BN-PAGE) further showed that the purified TOM complex formed a homogeneous assembly containing Tom40, Tom22, Tom20, Tom7, and Tom5 (Fig. 1B and Fig. S1). Owing to the lack of a suitable antibody, the presence of Tom6 could not be directly confirmed. The absence of Tom70 is consistent with its loose association with the TOM core complex reported previously (23,24). The estimated molecular mass of the TOM complex was approximately 530 kDa by gel filtration chromatography and approximately 400 kDa by BN-PAGE. These values were consistent with the predicted molecular mass of a trimeric TOM complex associated with detergent micelles, as suggested by previous biochemical studies (12,15).

**Figure 1.**
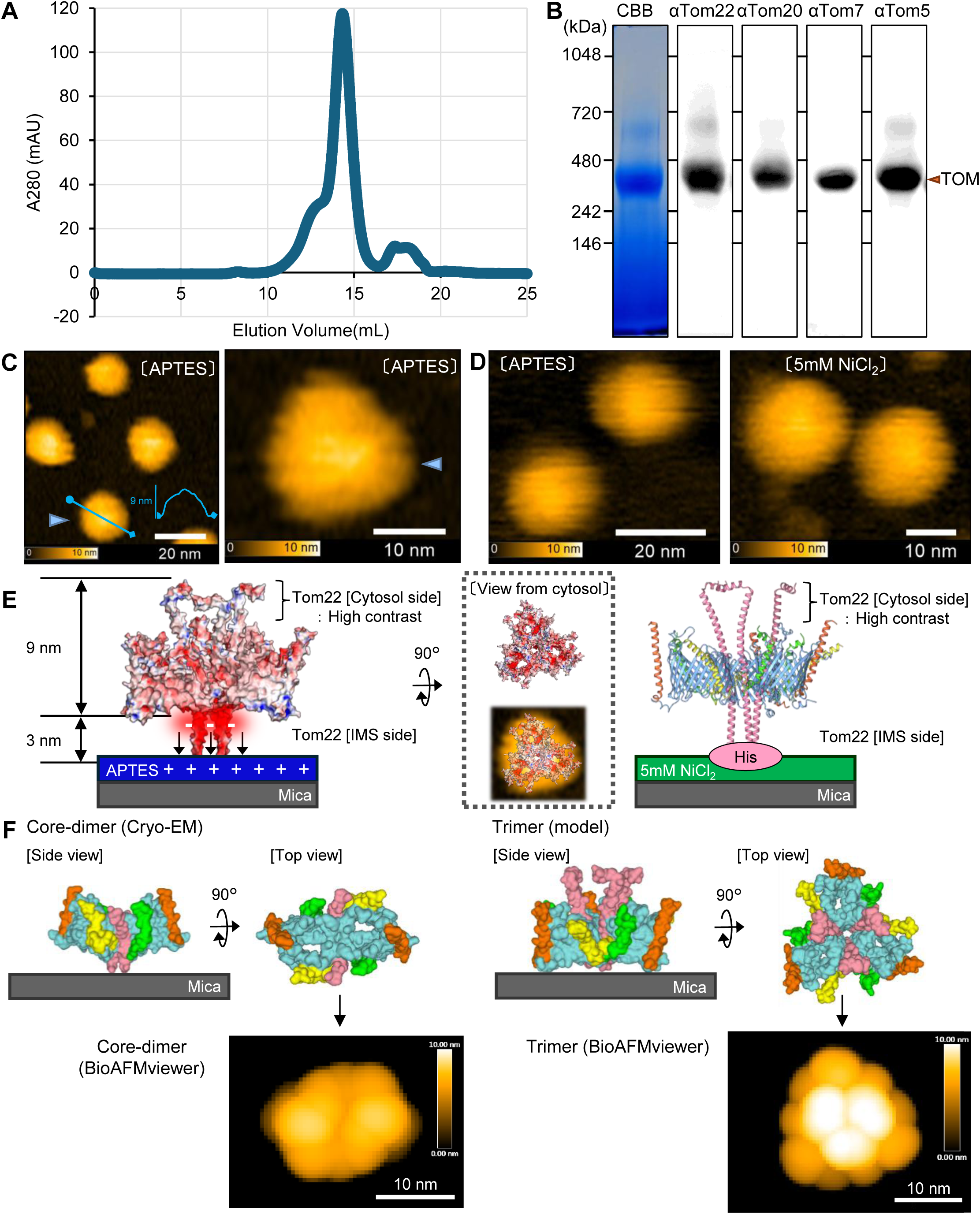
HS-AFM analysis of the purified TOM complex. (A) The size-exclusion chromatogram of the TOM complex under 0.01% GDN condition. (B) The largest peak of the size-exclusion chromatogram was analyzed by Blue Native-PAGE with CBB staining and western blot analysis. The purified TOM complex forms a 450 kDa protein complex containing Tom5, Tom7, Tom20 and Tom22. (C) HS-AFM images of the TOM complex on the APTES-mica surface. The wide range view was on the left and the close-up view was on the right. The blue arrow points to the same particle. Imaging conditions were either 2 frames per second with a scan size of 150 × 150 nm^2^ and a pixel resolution of 180 × 90 on the left panel and 20 frames per second with a scan size of 50 × 35 nm^2^ and a pixel resolution of 80 × 28 on the right panel. (D) Comparison between the TOM complexes on the APTES-mica and those on the NiCl2-mica surfaces. Imaging conditions were either 8.36 frames per second with a scan size of 60 × 60 nm^2^ and a pixel resolution of 160 × 80 on the left panel and 5 frames per second with a scan size of 60 × 42 nm^2^ and a pixel resolution of 80 × 56 on the right panel. The molecular images used in (C) and (D) were cropped from the original data. The uncropped images were included in the supplementary figures. (E) The schematic diagram using a trimeric structure model of the TOM complex. (left) The negative-charged patch of Tom22 on the IMS side is interacted with the positive-charged patch of the APTES-mica plate, and thus the cytosol side of the TOM complex was observed by HS-AFM. (right) the C-terminal His-tag of the Tom22 on the IMS side was fixed by Ni^2+^ on the mica surface. (F) The simulated HS-AFM images by BioAFMviewer. The simulated images were generated from the Cryo-EM structure of the core-dimer (left) and the model structure of the trimer (right).

HS-AFM imaging revealed that the purified TOM complexes predominantly exhibited homogeneous circular particles, consistent with a trimeric rather than a dimeric organization. (Fig. 1C). In the previously proposed trimeric model, three Tom40 channels are connected through Tom22 helices, resulting in clustering of the three Tom22 helices at the center of the complex (12). Tom22 is an important subunit which consists of an N-terminal cytosolic domain (∼100 amino acids), a transmembrane helix, and a short C-terminal IMS region (∼30 amino acids) (25,26). Because the mica surface was coated with 3-aminopropyltriethoxysilane (APTES), which imparts a positive charge and facilitates immobilization of TOM particles, the negatively charged IMS region of Tom22 was expected to interact with the mica surface (15,16) (Fig. 1E). Accordingly, the N-terminal cytosolic domain of Tom22 would be oriented toward the solvent and likely correspond to the bright central density observed in the HS-AFM images.

Based on the trimer structural model, the combined height of the transmembrane region and cytosolic domain is approximately 9 nm, whereas the IMS region of Tom22 extends ∼3 nm from the membrane (Fig. 1E, left panel). The TOM particles exhibited heights of approximately 8–10 nm by HS-AFM analysis (Fig. 1C), closely matching the expected dimensions of the transmembrane and cytosolic regions. These observations are consistent with a model in which the IMS helices of Tom22 are partially bent on the mica surface during HS-AFM imaging (Fig. 1E).

To further examine the orientation of the TOM complex on the mica surface, we additionally performed HS-AFM imaging using mica coated with nickel chloride to stabilize the His-tag located on the IMS side of Tom22 (Fig. 1D, Fig. 1E, right panel). Under these conditions, TOM particles exhibited structural features highly similar to those observed on APTES-coated mica, indicating that the TOM complexes were immobilized through their IMS regions in both conditions (Fig. 1D). These results support the interpretation that the current HS-AFM images represent a cytosolic view of the TOM complex (Fig. 1D). However, because immobilization through the His tag could potentially interfere with analyses of molecular dynamics, subsequent experiments were performed using TOM complexes attached to APTES-coated mica.

To further evaluate the observed HS-AFM structures, we generated simulated AFM images of both the TOM core dimer and trimer based on the cryo-EM structure and the trimeric model, respectively (Fig. 1F). The experimentally observed TOM particles displayed circular shapes with a bright central region, rather than the oval-shaped features predicted for the TOM core dimer, suggesting that the particles observed by HS-AFM more closely resembled a trimeric TOM complex (Fig. 1F).

### Dissociation of the trimeric TOM complex

HS-AFM observations revealed that the purified TOM complex was highly fragile. Because HS-AFM measures molecular height through repeated tapping with a nanometer-scale cantilever, continuous scanning gradually induced dissociation of the TOM complex during imaging (Fig. 2, Fig.S2 and Mov.1). We observed that individual circular TOM particles progressively transformed into triangular-shaped structures composed of three smaller lobes, which likely corresponded to three Tom40 channels. Notably, these triangular intermediates resembled particles previously reported in early electron microscopy studies (9,10).

**Figure 2.**
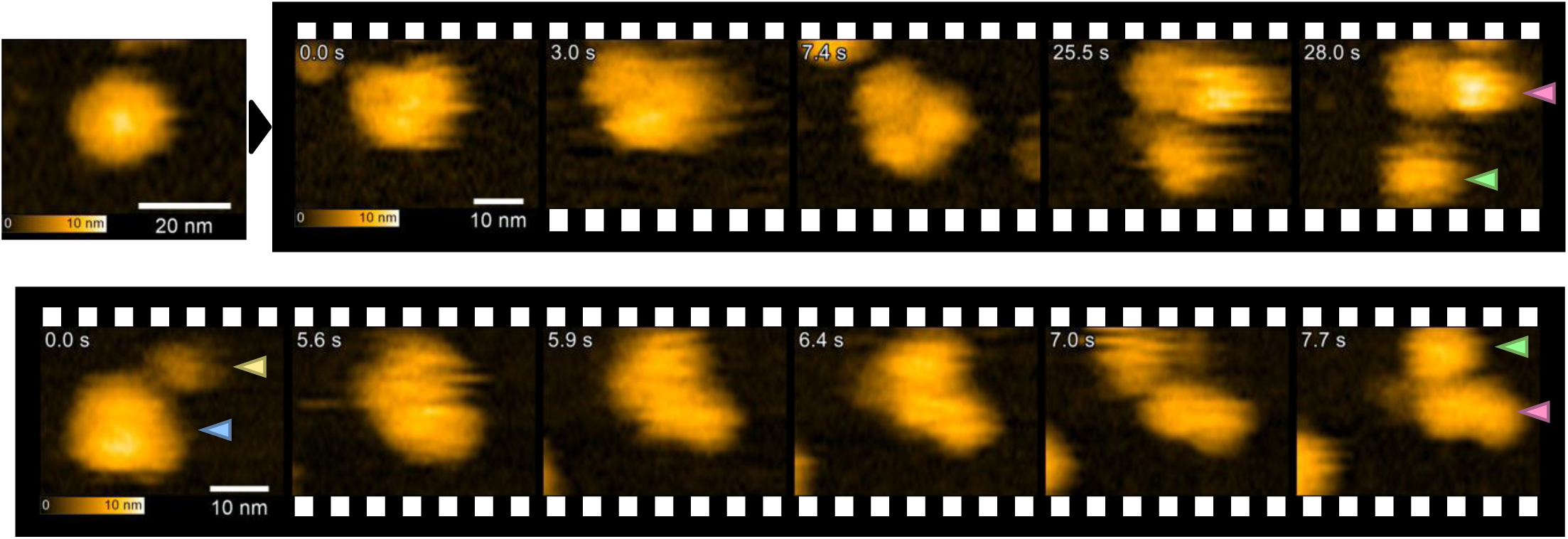
Dissociation of the TOM complex trimer. The trimeric TOM complex is dissociated into a dimer and a monomer with the passage of time. To demonstrate reproducibility, representative observations of two independent TOM complex particles were shown. The trimers, dimers, monomers and substrates are indicated by blue, red, green, and yellow arrows, respectively. Imaging conditions were either 7.68 frames per second with a scan size of 75 × 52 nm^2^ and a pixel resolution of 120 × 42 on the top panel and 6.67 frames per second with a scan size of 75 × 52 nm^2^ and a pixel resolution of 140 × 49 on the bottom panel. The molecular images used in this figure were also cropped from the original data. The uncropped images were included in the supplementary figures.

Subsequently, the trimeric TOM particles dissociated into a single isolated lobe and a pair of connected lobes, which likely corresponded to monomeric and dimeric TOM complexes, respectively (Fig.2, Fig.S2 and Mov.1). Because the dissociated particles were presumably re-surrounded by detergent micelles, reassociation of the monomeric and dimeric particles was rarely observed under the present experimental conditions. In addition, dissociation of the TOM trimer was further accelerated in the presence of substrate proteins or antibodies against the TOM complex, likely reflecting increased mechanical stress imposed on the TOM complex during HS-AFM imaging. Consistent with this intrinsic fragility, cryo-EM analysis of the same preparation preferentially resolved the TOM core-dimer rather than the trimeric assembly, indicating that the core-dimer constitutes a particularly stable configuration favored during cryo-EM grid preparation (Fig.S3). Taken together, these observations suggest that, although the trimeric TOM complex can be purified and maintained under detergent conditions, it remains structurally labile and readily transitions into smaller assemblies, including a stable core-dimer state.

### Biochemical identification of the trimeric TOM complex using three distinct tags

To further examine whether the purified TOM complex contains three Tom40 channels, we constructed a yeast strain simultaneously expressing three differently tagged Tom40 proteins fused to Strep-tag II, 3×FLAG, or PA tags, respectively. (Fig.3A) In this strain, Strep-tag II–, 3×FLAG-, and PA-tagged Tom40 proteins were coexpressed on the mitochondrial outer membrane (Fig. S4A). Assuming that these three tagged Tom40 proteins are incorporated into TOM complexes without strong assembly bias, TOM particles containing various combinations of tagged Tom40 subunits would be generated. Under a trimeric assembly model, TOM complexes simultaneously containing all three tags (“triple-tagged TOM complexes”) would be expected to occur at a frequency of 6/27 (approximately 22.2%), assuming equal probabilities for all 27 possible tag combinations.

**Figure 3.**
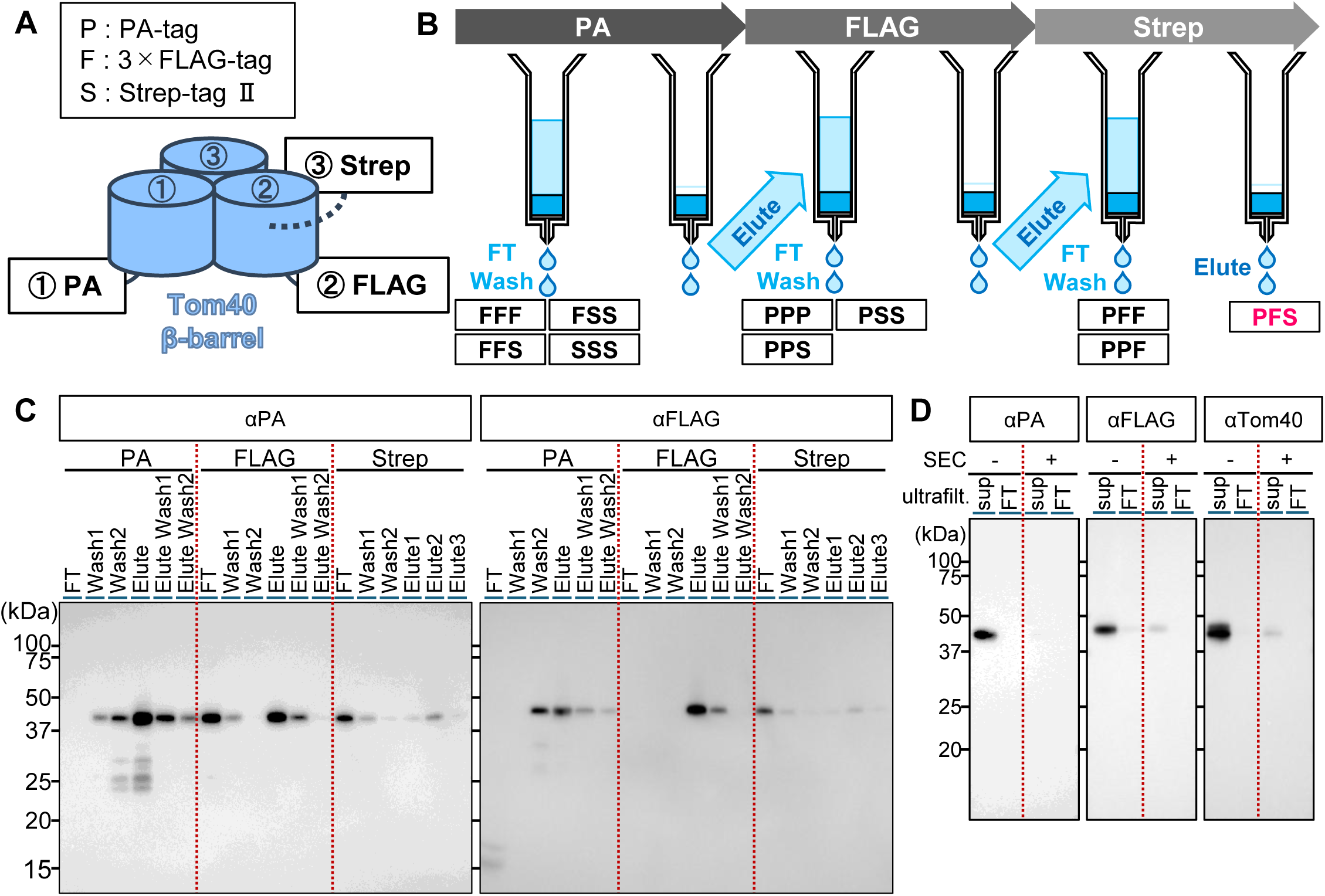
Identification of the trimeric TOM complex. (A) The schematic diagram of the triple-tagged TOM complex and the procedure of affinity purifications employing the PA, the 3×FLAG and the Strep II tags affinity resins. (B) The western blot analysis showing the three-step affinity purification process, with anti-PA and 3×FLAG tags antibodies and the HRP conjugated Strep-Tactin used for detection. (C, D) Gel filtration chromatogram of the triple tagged TOM complex and western blot analysis. Although the yield was low and no distinct peak could be detected, western blot analysis of the concentrated fractions around 14 mL, where the TOM complex was expected to elute, revealed a Tom40 band using anti- PA, 3×FLAG and Tom40 antibodies. Even though Tom40-Strep was not observed, likely because its concentration was below the detection limit of Strep-Tactin, it is believed that the purified TOM complex contains Tom40-Strep, as Strep II tag affinity purification was performed at the end of the three-step purification procedure.

Using isolated mitochondria, we sequentially purified TOM complexes through PA-, FLAG-, and Strep-tag affinity purification steps and analyzed the purified fractions by immunoblotting using anti-PA and anti-FLAG antibodies (Fig. 3B, 3C, Fig. S4B). Although the amount of purified material was below the detection limit of gel filtration chromatography, immunoblot analyses clearly detected PA- and 3×FLAG-tagged Tom40 proteins at the elution position corresponding to the TOM complex trimer (14–15 mL) (Fig. 3D, Fig. S4C). These results indicate that triple-tagged TOM complexes were successfully purified, albeit in small quantities. Together with the HS-AFM observations, the biochemical analyses support the interpretation that a substantial population of purified TOM complexes retains a trimeric organization after solubilization.

### Dynamic visualization of Tom20 receptors and substrate binding

To visualize substrate-binding regions within the purified TOM complex, we added dihydrofolate reductase (DHFR) fused to the presequence of *Neurospora crassa* ATP synthase subunit 9 (pSu9) during HS-AFM observation of the TOM complex (12,27). Although pSu9-DHFR particles rapidly diffused across the mica surface, they also transiently interacted with TOM complex particles (Fig. 4A). Real-time HS-AFM imaging demonstrated that one to three pSu9-DHFR particles could simultaneously associate with a single TOM complex, suggesting that individual channels within the TOM complex possess distinct substrate-interaction regions.

**Figure 4.**
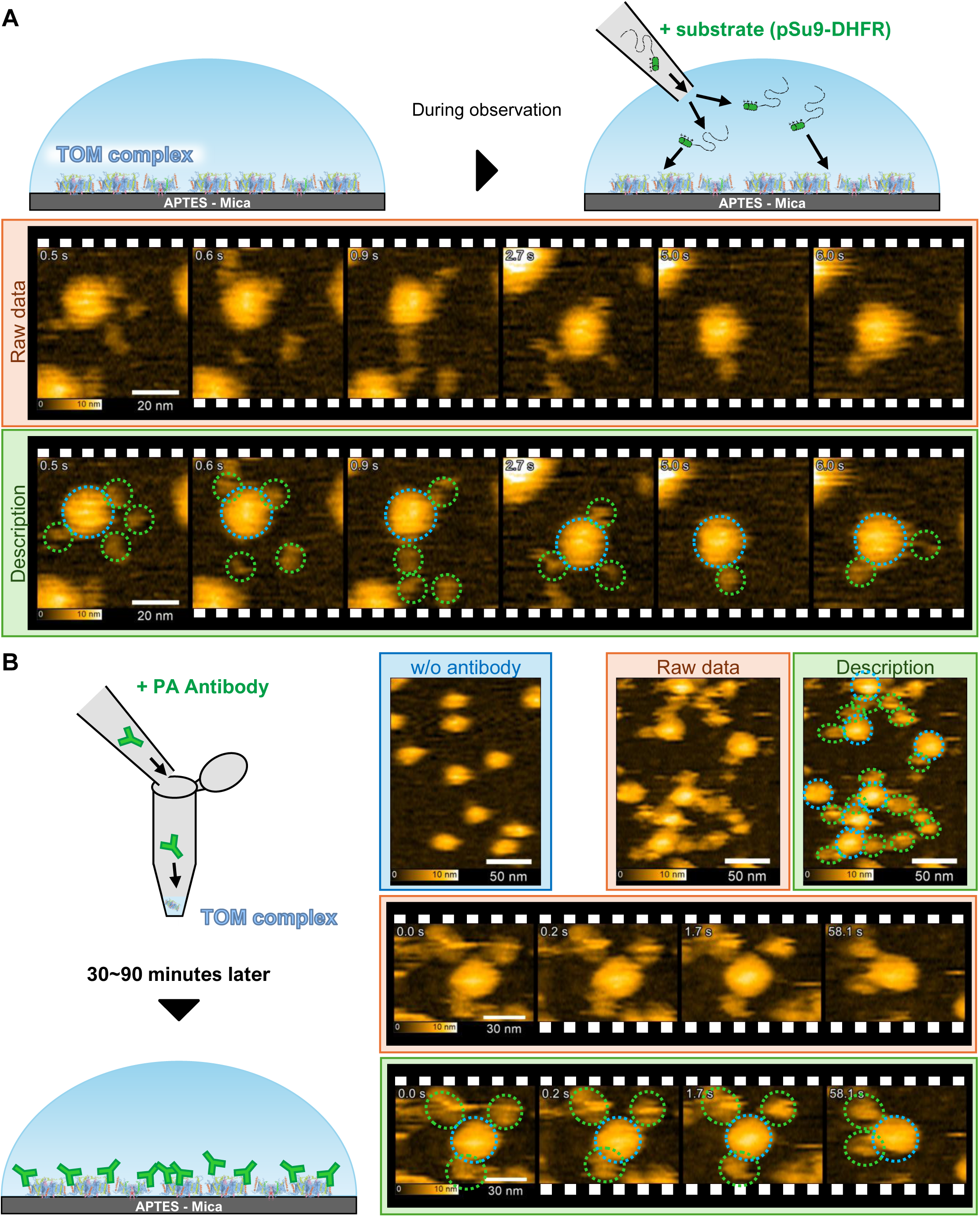
Real-time observation of the position of the Tom20 receptor subunit and its substrate recognition. (A) Schematic diagram of the substrate addition experiment was briefly described above. The sequential images of the TOM complex and pSu9-DHFR were shown. The upper panel indicated only the raw data, while the bottom panel included descriptions of each individual particle. The images in the upper and bottom panels are identical. The anti-PA antibodies interacted with the TOM complex, which suggests the spatial positions of the Tom20 subunits of the TOM complex in the HS-AFM images. The substrate protein pSu9-DHFR and the TOM complexes are surrounded by green and blue circles, respectively. Imaging conditions were either 6.68 frames per second with a scan size of 80 × 80 nm^2^ and a pixel resolution of 120 × 60. (B) Schematic diagram of the anti-PA tag antibody addition experiment was briefly described on the left side. The representative images of the crosslinked-TOM complex were shown separately in the absence and presence of antibodies. The sequential images of the cross-linked TOM complex with anti-PA tag antibodies were shown. The upper panel indicated only the raw data, while the bottom panel included descriptions of each individual particle. The images in the upper and bottom panels are identical. The anti-PA antibodies and the TOM complexes are surrounded by green and blue circles, respectively. Imaging conditions were either 1.05 frames per second with a scan size of 300 × 210 nm^2^ and a pixel resolution of 400 × 70 on the top panel and 5.01 frames per second with a scan size of 100 × 70 nm^2^ and a pixel resolution of 200 × 35 on the bottom panel. The molecular images used in this figure were also cropped from the original data. The uncropped images were included in the supplementary figures.

Previous electron microscopy studies suggested that the receptor subunit Tom20 is localized at the outer peripheral region of the trimeric TOM complex and functions in precursor recognition (11). We therefore attempted to visualize the position of Tom20 in HS-AFM images and compare its localization with the substrate-binding behavior observed above. To this end, we purified a PA-tagged TOM complex in which a PA tag was fused to the C-terminal cytosolic region of Tom20 and incubated the purified complex with anti-PA tag antibodies. If Tom20 subunits are localized at the periphery of the TOM complex, one to three antibody particles would be expected to associate with individual TOM complexes.

As described above, TOM complex trimers were highly fragile and frequently dissociated during HS-AFM observation. Addition of antibodies further destabilized the TOM complexes during imaging, thereby interfering with stable visualization (Fig. S2). To overcome this limitation, we employed the GraFix method, which combines glutaraldehyde cross-linking with sucrose density gradient centrifugation, to stabilize TOM complex particles before HS-AFM analysis (28) (Fig. S5).

Prior to the antibody-binding experiments, we evaluated the efficiency of GraFix-mediated cross-linking. SDS-PAGE analysis revealed broad high-molecular-weight signals, indicating that TOM complex subunits were cross-linked by glutaraldehyde (Fig. S5AB). In addition, BN-PAGE analysis showed that the cross-linked TOM complex migrated at a position similar to that of the untreated complex, suggesting that glutaraldehyde predominantly cross-linked subunits within the TOM complex without forming larger aggregates (Fig. S5CD).

When the cross-linked TOM complex was incubated with anti-PA tag antibodies and analyzed by HS-AFM, we successfully captured molecular images showing antibody particles associated with TOM complexes (Fig. 4B). One to three antibody particles interacted with individual TOM complexes, suggesting that Tom20 subunits are localized at peripheral regions of the trimeric TOM complex (Fig. 4B).

Because the GraFix method successfully stabilized TOM complex trimers for HS-AFM observation, we next attempted structural analysis by single-particle cryo-EM. Although this analysis again yielded predominantly a dimeric structure resembling those reported previously (15, 16), the 2D class averages also contained a putative trimeric form corresponding to the three Tom40 pores. (Fig. S5E). Despite the presence of this trimeric species, we were unable to obtain a high-resolution 3D reconstruction of the trimeric complex. These observations suggest that trimeric TOM complexes, although likely present on the cryo-EM grids, may not adopt a sufficiently stable or uniform conformation within detergent micelles suitable for high-resolution single-particle reconstruction, such that their structural heterogeneity or conformational flexibility limited efficient alignment and high-resolution reconstruction during image processing. In this regard, our observations are consistent with previous negative-staining electron microscopy studies that identified trimeric TOM particles at the single-particle level (9,10).

## Discussion

Since the core-dimer structure of the TOM complex from *Neurospora crassa* was determined at 6.8 Å resolution in 2017 (14), high-resolution TOM complex structures at near-atomic resolution have been reported by several groups (15–22). Nevertheless, cryo-EM structural analyses have predominantly captured the TOM complex in its core-dimer form, whereas tetrameric and hexameric assemblies have occasionally been observed (16,18). Notably, the tetrameric structure essentially represents a dimerization of two TOM core dimers. In contrast, the precise structure corresponding to the trimeric TOM complex proposed in earlier studies has not yet been determined, despite previous electron microscopy studies suggesting the presence of trimeric TOM particles (9–12). Previously, we also observed trimeric TOM particles by HS-AFM during cryo-EM structural studies but did not perform detailed characterization (15). In the present study, we further analyzed the trimeric TOM complex by HS-AFM to investigate its structural features and dynamic behavior.

Based on the present observations, we propose a possible relationship between the trimeric TOM complex and the TOM core dimer. As shown in Fig. 2, trimeric TOM particles frequently dissociated into dimeric and monomeric particles during HS-AFM observation. The dimeric particles observed by HS-AFM likely correspond to the TOM core-dimer structures previously identified by cryo-EM analysis (Fig. 2). This interpretation is further supported by the similarity between the experimentally observed HS-AFM images and the simulated ones generated from the reported TOM core-dimer structures (Fig. 1F, Fig. 2). During cryo-EM grid preparation and image processing, one Tom40 molecule within the trimeric assembly may become structurally unstable or partially dissociated, thereby favoring formation of the TOM core dimer (Fig.S3).

However, this possibility does not imply that the TOM core dimer is an experimental artifact. Because previous biochemical and structural analyses based on the TOM core dimer successfully explained key aspects of mitochondrial protein translocation, the TOM core dimer likely represents a functional and energetically stable structural state on the mitochondrial outer membrane (15). It is therefore conceivable that monomeric TOM particles are associated with the stable core-dimer unit to form a trimeric assembly (Fig. 2). In this model, the TOM trimer may adopt an asymmetric organization, in which a stable core dimer serves as the central structural unit and an additional monomer associates more dynamically with the complex, rather than three completely equivalent channels functioning in a fully symmetric manner. Although detailed structural analysis of the TOM trimer will require future high-resolution studies, the current findings support the existence of trimeric TOM assemblies while suggesting that the energetically stable core-dimer region is preferentially resolved by cryo-EM analysis. We hope that the present study will facilitate future structural determination of the TOM trimeric complex.

Within the TOM complex, Tom70, Tom22, and Tom20 function as receptor subunits involved in substrate recognition (29,30). Because Tom70 dissociated during solubilization of yeast mitochondria, its functional contribution could not be evaluated in the present HS-AFM analyses (14). Tom22 additionally functions as a structural linker between Tom40 channels and is positioned at the center of the proposed trimeric TOM complex model (12). Even in the TOM core-dimer structures, two Tom22 helices are located between adjacent Tom40 channels, serving as a scaffold essential for maintaining TOM complex integrity (15,16).

In contrast, Tom20 is known to interact relatively weakly with the TOM complex and is thought to move with some degree of flexibility around the periphery of the complex (12,15). Although Tom20 is frequently disordered in TOM core-dimer structures, several cryo-EM studies have identified its approximate position by stabilizing the TOM complex through cross-linking or substrate addition (19–22). Owing to its structural flexibility and the variable positioning of its transmembrane helix, the precise binding site of Tom20 has remained difficult to define. Consequently, local structural resolution in this region is often limited, and atomic modeling of Tom20 is frequently incomplete. In the previously reported TOM core-dimer structures, the receptor domains of Tom20 are positioned above the Tom40 channels (20,21). By contrast, earlier moderate-resolution electron microscopy and cross-linking studies of the TOM trimer suggested that Tom20 localizes to the peripheral region of the triangular trimeric assembly (11,12).

In the present HS-AFM analyses, we tracked the molecular behavior of pSu9-DHFR in the presence of the TOM complex and observed that one to three pSu9-DHFR molecules associated with peripheral regions of individual TOM complexes (Fig. 4A). Because previous structural and biochemical studies suggested that Tom20 localizes to the periphery of the trimeric TOM complex, it is reasonable to interpret these peripheral substrate interactions as reflecting binding to Tom20 (11). In contrast, no clear substrate interaction was observed at the central region of the TOM complex, where Tom22 is predicted to accumulate. This observation may reflect the distinct role of Tom22 as a structural organizer of the TOM complex rather than a primary substrate receptor and is consistent with the idea that substrate recognition primarily depends on Tom20 or Tom70 (31). Furthermore, HS-AFM visualization of interactions between anti-PA tag antibodies and peripheral regions of the PA-tagged TOM complex further supported the peripheral localization of Tom20 and its involvement in substrate recognition (Fig. 4B). Consistent with this interpretation, previous cryo-EM analyses of Tom20-containing TOM complexes identified a long α-helical segment between the receptor domain and the transmembrane helix, suggesting that the position of Tom20 can be flexibly adjusted relative to the TOM complex (21,22).

Thus, through HS-AFM analysis of the TOM complex, we supported that the TOM complex predominantly adopts a trimeric organization while also existing in stable dimeric and monomeric states. Furthermore, our analyses provided insight into the spatial arrangement of the Tom20 receptor, consistent with previous models of the trimeric TOM complex. While recent cryo-electron microscopy studies have substantially advanced our understanding of the TOM core-dimer structure, the present single-molecule analyses enabled characterization of the trimeric TOM complex, which has not yet been fully resolved by cryo-EM analysis.

Whether the TOM complex physiologically requires a trimeric architecture and what functional advantages are conferred by the presence of three channels remain important open questions. Nevertheless, the present study enabled characterization of the trimeric TOM complex *in vitro* and provides a foundation for future structural and functional analyses of TOM complex assembly and dynamics.

## Methods

### Yeast strain, plasmids and primers

To construct the *S. cerevisiae* strain for overexpression of the TOM complex, we introduced the *GAL1* promoter using the Yeast Tool Kit (YTK) and Golden Gate assembly, following a strategy similar to that used in a recent structural study (16,32,33) (Supplementary Table 1). The coding sequences (CDSs) of Tom40, Tom22, Tom20, Tom7, and Tom5 were amplified by PCR, whereas the Tom6 gene was chemically synthesized. Each gene was individually inserted into the pYTK001 entry plasmid (Supplementary Table 2). To enable affinity purification of the TOM complex, a Strep-tagⅡsequence (WSHPQFEK) with a GG linker was fused to the C-terminus of Tom40. In addition, a 10×His tag with a GG linker was fused to the C-terminus of Tom22 to facilitate immobilization of the TOM complex on mica surfaces during HS-AFM analysis. (see the HS-AFM observation section below). The *GAL1* promoter (YTK030), the CDS of each TOM complex subunit, and the *ENO1* terminator (YTK061) were assembled into the pYTK095 cassette plasmid. The resulting cassette plasmids encoding individual TOM complex subunits were subsequently inserted into the pYTK096 plasmid containing the *URA3* auxotrophic marker (Supplementary Table 3). The final vector was linearized with NotI and introduced into the W303 yeast strain using a standard lithium acetate transformation method. Transformants were selected on SCD-ura medium containing 0.67% (w/v) yeast nitrogen base without amino acids, 0.5% (w/v) casamino acids, 2% (w/v) glucose, 0.002% (w/v) adenine sulfate, and 0.002% (w/v) L-tryptophan.

To generate the strain expressing triple-tagged Tom40 proteins, two additional Tom40 constructs fused to either a PA tag or a 3×FLAG tag through GG linkers were subcloned into the pYTK095 vector. These tagged Tom40 constructs were tandemly integrated into a pYTK096-based vector containing the LEU2 auxotrophic marker (Supplementary Table 3). The TOM complex overexpression strain described above was subsequently transformed with this vector and cultured in SCD-leu-ura medium containing 0.67% (w/v) yeast nitrogen base without amino acids, 1.74% (w/v) SC-leu-ura supplement mixture (MP Biomedicals), and 2% (w/v) glucose. The resulting yeast strain simultaneously overexpressed three Tom40 variants fused to PA, 3×FLAG, and Strep-tag II sequences, respectively.

For HS-AFM visualization of antibody binding, we additionally constructed another TOM complex overexpression strain in which a PA tag was fused to the C terminus of Tom20, together with the Strep-tag II fused to Tom40 and the 10×His tag fused to Tom22.

### Protein expression and purification

Yeast strains were cultured in YPD (1%(w/v)yeast extract, 2%(w/v)polypeptone, 0.1%(w/v)glucose)medium until OD₆₀₀ reaches 0.6, then galactose was added to induce the transcription by the *GAL1* promoter, and cultured in 4l YPGal(1%(w/v)yeast extract, 2%(w/v)polypeptone, 2%(w/v) galactose) medium at 30°C for 20 hours. The following procedure for mitochondrial isolation was performed in accordance with the recent publication (33). In brief, the yeast cells were collected and resuspended with Tris-SO4 buffer (100 mM Tris SO4 [pH 9.4], 10 mM DTT). After incubating with gentle shaking at 90 rpm and 30 °C for 10 min, the cell pellet was collected and incubated in sorbitol buffer (1.2 M sorbitol, 20 mM Tris-HCl [pH 7.5]) containing zymolyase 20T (zymolyase 5 mg/yeast 1 g) with shaking at 135 rpm and 30 °C for 30 min. After washing out the zymolyase with sorbitol buffer, cell pellet was disrupted in breaking buffer (600 mM sorbitol, 10 mM Tris-HCl [pH 7.5], 1 mM EDTA) using a tight-fitting Dounce homogenizer. Then, mitochondrial membrane fraction was isolated through multiple centrifugations and washed by SEM buffer (250 mM sucrose, 1 mM EDTA, 10 mM MOPS·NaOH [pH 7.2]).

Isolated mitochondria were solubilized by solubilization buffer (20 mM Tris-HCl [pH 7.5], 150 mM NaCl, 1% digitonin) at 4 °C overnight, and centrifuged at 20,000 g at 4 °C for 20 min. The supernatant was mixed with 1mL of Strep-Tactin® 4Flow® high capacity resin (IBA) and incubated at 4 °C for more than 4 hours. The resin was washed with Affinity buffer (50 mM Tris-HCl pH7.5, 150 mM NaCl, 0.1% GDN), and the TOM complex was eluted by Affinity Buffer containing 3 mM D-desthiobiotin.

The eluted TOM complex was concentrated using an Amicon Ultra (Merck Millipore, 100 kDa MW cut-off) ultrafiltration kit by centrifugation at 5,000 g (final volume was approximately 500 μL). The TOM complex was further purified by gel filtration chromatography using a Superose 6 Increase 10/300 GL column (cytiva) equilibrated with the SEC buffer (20 mM Tris [pH 7.5], 150 mM NaCl, 0.01% GDN).

Especially for Cryo-EM analysis, the overexpression strain was cultured in 36-40 L of YPD medium. The purification procedures were almost same as those for HS-AFM analysis, but with 4 mL of Strep-Tactin® 4Flow® high capacity resin. To obtain high yield, even the TOM complex that passed through the column was recovered from flowthrough fraction.

### Purification of the triple-tagged TOM complex

The yeast strain overexpressing the triple-tagged TOM complex was cultured in 18L of YPD medium. Mitochondrial membrane fraction was isolated and solubilized in the same way as described above. Then, the triple-tagged TOM complex was sequentially purified using three affinity resins. First, the supernatant was mixed with 1ml of Anti PA tag Antibody Beads (FUJIFILM) and incubated at 4℃ for 2 hours. The resin was washed with Affinity buffer and incubated in Affinity Buffer containing 0.1 mg/ml of PA tag Peptide (FUJIFILM) for 1 hour. After that, the eluate TOM complex was collected. Furthermore, the resin was washed by 10mL of Affinity buffer without any PA peptide (elute wash). The elution and elute wash fractions were mixed and subjected to the 400 μL of ANTI-FLAG® M2 Affinity Gel (Merck) as a second affinity purification. After incubating at 4℃ for 1 hour, the resin was washed with Affinity buffer and incubated in Affinity Buffer containing 0.1 mg/ml of 3×FLAG Peptide (ProteinArk) for 1 hour. After that, the elution and elute wash fractions were collected and subjected to 1 ml of Strep-Tactin® 4Flow® high capacity resin (IBA) as a third affinity purification. After incubating at 4℃ overnight, the resin was washed with Affinity buffer, and the triple-tagged TOM complex was eluted by Affinity Buffer containing 3 mM D-desthiobiotin.

The eluted triple-tagged TOM complex was concentrated using an Amicon Ultra (Merck Millipore, 100 kDa MW cut-off) by centrifugation at 5,000 g (final volume was approximately 500 μL). The triple-tagged TOM complex was analyzed by gel filtration chromatography using a Superose 6 Increase 10/300 GL column (cytiva) equilibrated with the SEC buffer. The elution fractions around 14-15mL were mixed and concentrated Amicon Ultra (Merck Millipore, 100 kDa MW cut-off) and analyzed by SDS-PAGE.

### Grafix

To prevent dissociation, we applied the gradient fixation method (GraFix) to the structural analysis (28). First, the TOM complex was purified by Strep-tag affinity purification using 50 mM HEPES-KOH pH7.4 instead of Tris-HCl buffer and gel filtration chromatography and concentrated by Amicon Ultra (100 kDa MW cut-off) up to 400 μL volume. Then, a 17-steps sucrose gradient was prepared manually, in which each fraction volume was 400 μL (Supplementary Table 4). The low-density buffer was composed of 20 mM HEPES-KOH pH7.4, 150 mM NaCl, 0.01 % GDN, and 5 % (w/v) sucrose, while the high-density buffer contained 20 mM HEPES-KOH pH7.4, 150 mM NaCl, 0.01 % GDN, 45 % (w/v) sucrose, and 0.8 % glutaraldehyde. 200 μL of the purified TOM complex was applied on the top. Centrifugation was carried out at 100,000 g (Accel:9, Decel:1) at 4°C for 13 hours using a S50ST rotor (Himac).

Crosslinking efficiency was evaluated by SDS-PAGE with silver staining. The fractions between 4 to 9 were collected and dialyzed using Tube-O-DIALYZER (G-biosciences) in 200 mL of SEC buffer at 4°C. After three hours of dialysis, the buffer was replaced, and dialysis was continued overnight. For cryo-EM analysis the dialyzed TOM complex was concentrated by Amicon Ultra (100 kDa MW cut-off) up to 30 μL (Final concentration: 18.8 mg/mL).

### Immunoblotting

The gel was transferred onto a Transblot Turbo Pack membrane (PVDF). Rabbit anti-Tom40, Tom22, Tom20, Tom7 and Tom5 primary antibodies (1:1,000-10,000) were used to evaluate the TOM complex formation in each purification. Rat anti-PA and mouse anti-FLAG antibodies (1:5,000, 1:10,000, respectively) were used to evaluate the PA-tagged TOM complex and the triple-tagged TOM complex. Detection was carried out using Horseradish peroxidase (HRP)-conjugated goat anti-rabbit, rat, mouse antibodies (1:5000). HRP-conjugated Strep-Tactin (IBA) was also used to evaluate the Strep II tag that fused to Tom40 in all the TOM complex purified in this study. Signals were detected with a LuminoGraph I (ATTO) using chemiluminescence.

### HS-AFM Observation

The purified sample was collected only from the apex of the gel filtration chromatogram and diluted with an imaging buffer containing 10 mM Tris-HCl, pH 7.5, 30 mM NaCl, 10% glycerol and 0.01% GDN for dispersed observation. HS-AFM analysis was performed in tapping mode at room temperature using a laboratory-built HS-AFM setup (34–36). We used short cantilevers (BL-AC7DS-A2, Olympus; spring constant, ∼0.25 N/m; resonant frequency and quality factor in liquid, ∼800 kHz and ∼1.5, respectively). The probe tip was grown on the original tip end of a cantilever through electron beam deposition using ferrocene and was further sharpened using a radio-frequency plasma etcher (Tergeo, PIE Scientific LLC) under an argon gas atmosphere (Direct mode, 10 sccm, and 20 W for 1.5 min). To increase spatial sampling, we employed non-square pixels in all observations; the pixel size along the fast axis (X) was set to 2∼4× smaller than that along the slow axis (Y) (37). The general procedure for HS-AFM observation of protein molecules was described previously (38). In brief, a glass sample stage (diameter, 2 mm; height, 2 mm) with a thin mica disc (1.5 mm in diameter and 0.05 mm thick) glued to the top by epoxy was attached onto the top of the Z-scanner by a drop of nail polish. The mica surface was treated by (3-aminopropyl)triethoxysilane (APTES) diluted to 1:10,000 or 5 mM nickel chloride to facilitate immobilizing of the TOM complex particles on to the mica surface. Especially for the TOM complex crosslinked by GraFix method, APTES was diluted to 1:2,000 due to higher mobility of the TOM complex particle on mica surface. A drop (2 μL) of the purified TOM complex was deposited onto an APTES-treated mica surface and incubated for 3 min to allow partial adsorption. The surface was gently rinsed with the imaging buffer to remove unbound molecules, and then molecular dynamics were measured in a liquid cell containing ∼60 μl of the same buffer. The data collection was conducted using UMEX, custom-built software.

In the analysis of the interaction between the substrate protein pSu9-DHFR and the TOM complex, 5 μL of substrate was added to the buffer chamber twice during observation of the TOM complex. After a fer minutes, pSu9-DHFR gradually began to appear on the mica surface, and its molecular behavior was analyzed. For tracking the interaction between the anti-PA-tag antibodies and the PA-tagged TOM complex, we added one-tenth the amount of antibody (Anti PA tag, Rat Monoclonal Antibody (FUJIFILM)) to the cross-linked PA-tagged TOM complex. After incubating for 30–90 minutes, HS-AFM analysis was performed.

Simulated AFM images were analyzed by the BioAFMviewer (39) with a probe curvature radius of 10 nm, a cone angle of 10 degrees, and a pixel size of 1 nm.

### Cryo-EM analysis and data processing

The TOM complex without cross-link was concentrated up to 16.5 mg/mL and loaded onto the Chameleon system (SPT Labtech). A total of 40 nl of the sample solution was dispensed to the glow-discharged Quantifoil Active Au 300-mesh R1.2/2 grids with Cu nanowires (SPT Labtech) and the grids were plunged into liquid ethane and frozen in liquid nitrogen. The cross-linked TOM complex sample was concentrated up to 10.9 mg/mL and loaded onto a glow-discharged Quantifoil R1.2/1.3, Au,300 mesh grids. The grids were blotted by Vitrobot Mark IV (Thermo Fisher Scientific) and plunge-frozen in liquid ethane.

Micrographs for all datasets were collected with a Titan Krios G3i microscope (Thermo Fisher Scientific) running at 300 kV and equipped with a Gatan Quantum-LS Energy Filter (GIF) and a Gatan K3 Summit direct electron detector in the electron counting mode (The University of Tokyo, Japan). Datasets were collected with a total dose of approximately 50 electrons per Å^2^ per 48 frames by the standard mode, using the EPU software (Thermo Fisher Scientific). The dose-fractionated movies were subjected to beam-induced motion correction and dose weighting using Patch Motion Correction, and the contrast transfer function (CTF) parameters were estimated using Patch-based CTF estimation in cryoSPARC v4.4.0 (40)

Details of data collection and image processing are described in Supplementary Table 5. Cryo-EM data were processed by using cryoSPARC v4.4.0 (40). For the TOM complex without cross-link, 4,987,912 particles were initially picked from the 8,397 motion-corrected and dose-weighted micrographs using Template picker, followed by several rounds of reference-free 2D classification to curate particle sets. The particles were further curated by heterogeneous refinement, using maps derived from ab initio reconstruction as templates. The resulting 290,817 particles were subjected to non-uniform refinement, yielding a map at overall resolutions of 3.57 Å according to the FSC criterion of 0.143 (41,42). Molecular graphics and cryo-EM density map figures were prepared using CueMol (http://www.cuemol.org) and UCSF ChimeraX (43). The cryo-EM density map has been deposited in the Electron Microscopy Data Bank under accession code EMD-81859.

## Acknowledgements

The authors express their gratitude to Araiso lab members for their valuable discussion, Ms. Kana Takata for technical assistances and Dr. Akihiro Inazu for supporting in setting up the research environment. Also, the authors thank Dr. Toshio Ando, Ms. Aimi Makino, Ms. Risa Omura and Ms. Kayo Nakatani for technical support of the HS-AFM experiments. This work was supported by JSPS KAKENHI to Y.A. (20H02583, 22K19268 and 23H01814), JST FOREST program (JPMJFR200F) to Y.A. and JST SPRING program (JPMJSP2135) to Na.K.. Y.A. was also supported by Leading Initiative for Excellent Young Researchers, MEXT, Japan. The following grants are also acknowledged: Platform Project for Supporting Drug Discovery and Life Science Research (Basis for Supporting Innovative Drug Discovery and Life Science Research (BINDS)) from AMED under grant numbers JP25am121002 (support number 3272) (to O.N.) and JP26ama121029j0005 (support number 4859) (to K.I.); Hokuriku Bank Research Grant for Young Scientists (to Y.A.); The Mitsubishi Foundation Research Grants in the Natural Science (to Y.A.); and Naito Memorial Research Grant for Supporting the Development of the Next Generation (to Y.A.). This work was partly supported by Bio-SPM Collaborative Research Project, WPI-Nano Life Science Institute, Kanazawa University (to Y.A.). AI was used for language polishing.

## Author contributions

Na.K. performed most of the experiments such as sample preparation, biochemical analyses and HS-AFM analysis. K.K. and S.K. also performed sample preparation. S.N.O. performed and O.N. supervised cryo-EM measurement and data processing. K.I. performed 3D structural model building of the TOM trimeric complex. H.I., K.U., R.A. and No.K. supported HS-AFM data collection and analysis. Y.A. supervised the project and performed HS-AFM analysis. Na.K., T.E., No.K. and Y.A. designed research. Na. K., S.N.O. and Y.A. wrote the paper. No.K. and T.E. reviewed and edited the paper. All authors discussed the results of the experiments and commented on the manuscript.

## Supplemental Information

### Supplementary Figure Legends

**Figure S1.**
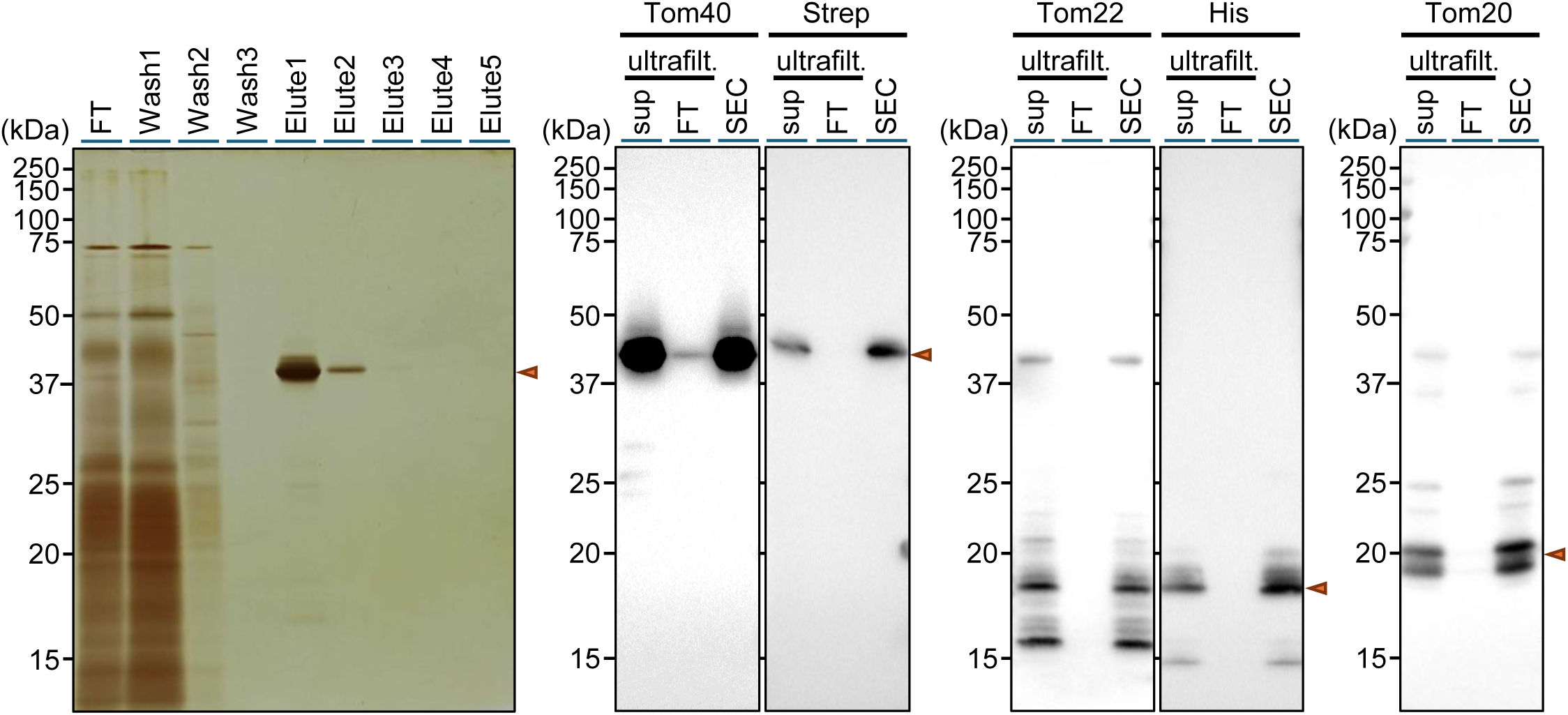
Purification of the TOM complex. The SDS-PAGE gel with silver staining *(left)* and western blot analyses *(right)*.

**Figure S2.**
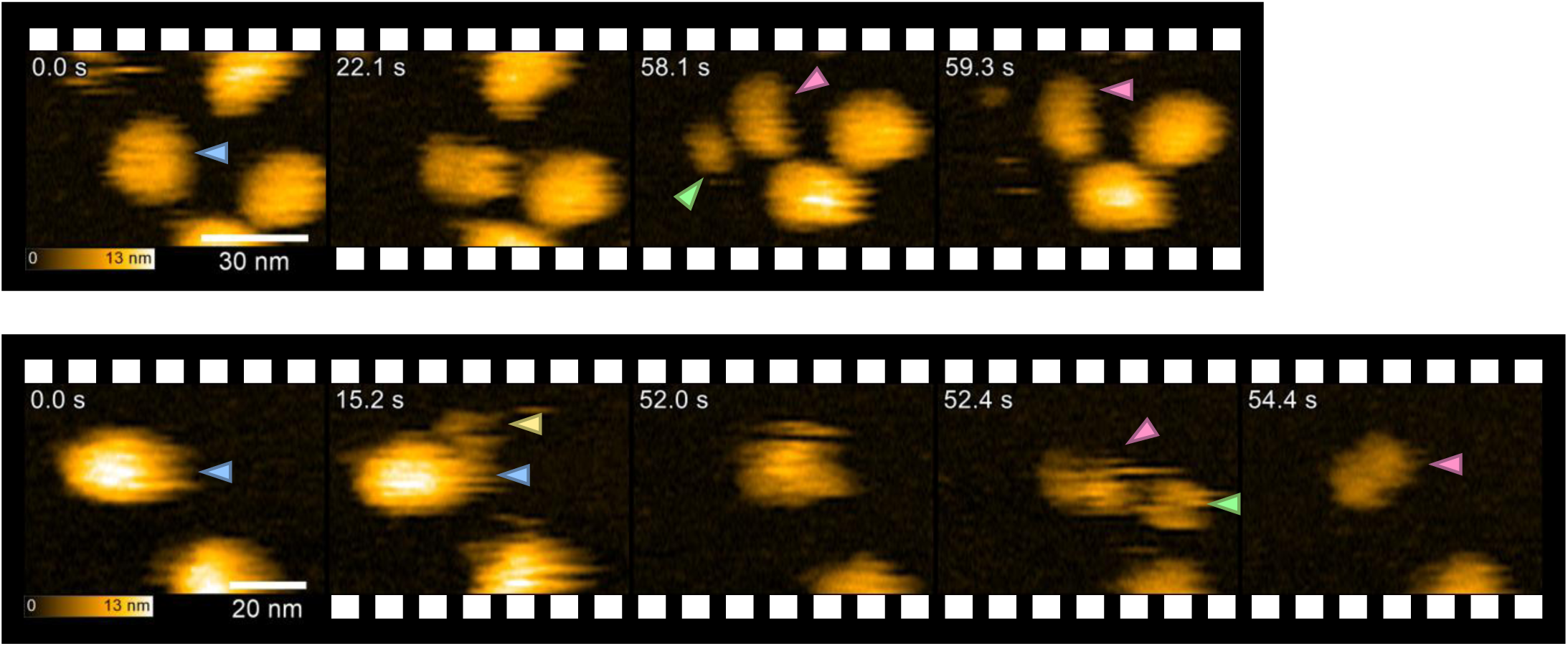
Cryo-EM analysis of the TOM complex without crosslinking. (A) Representative cryo-EM micrograph, recorded on a 300-kV Titan Krios microscope with a K3 camera. (B) Representative 2D class average images. (C) Final cryo-EM 3D reconstruction of the same TOM complex sample used for HS-AFM.

**Figure S3.**
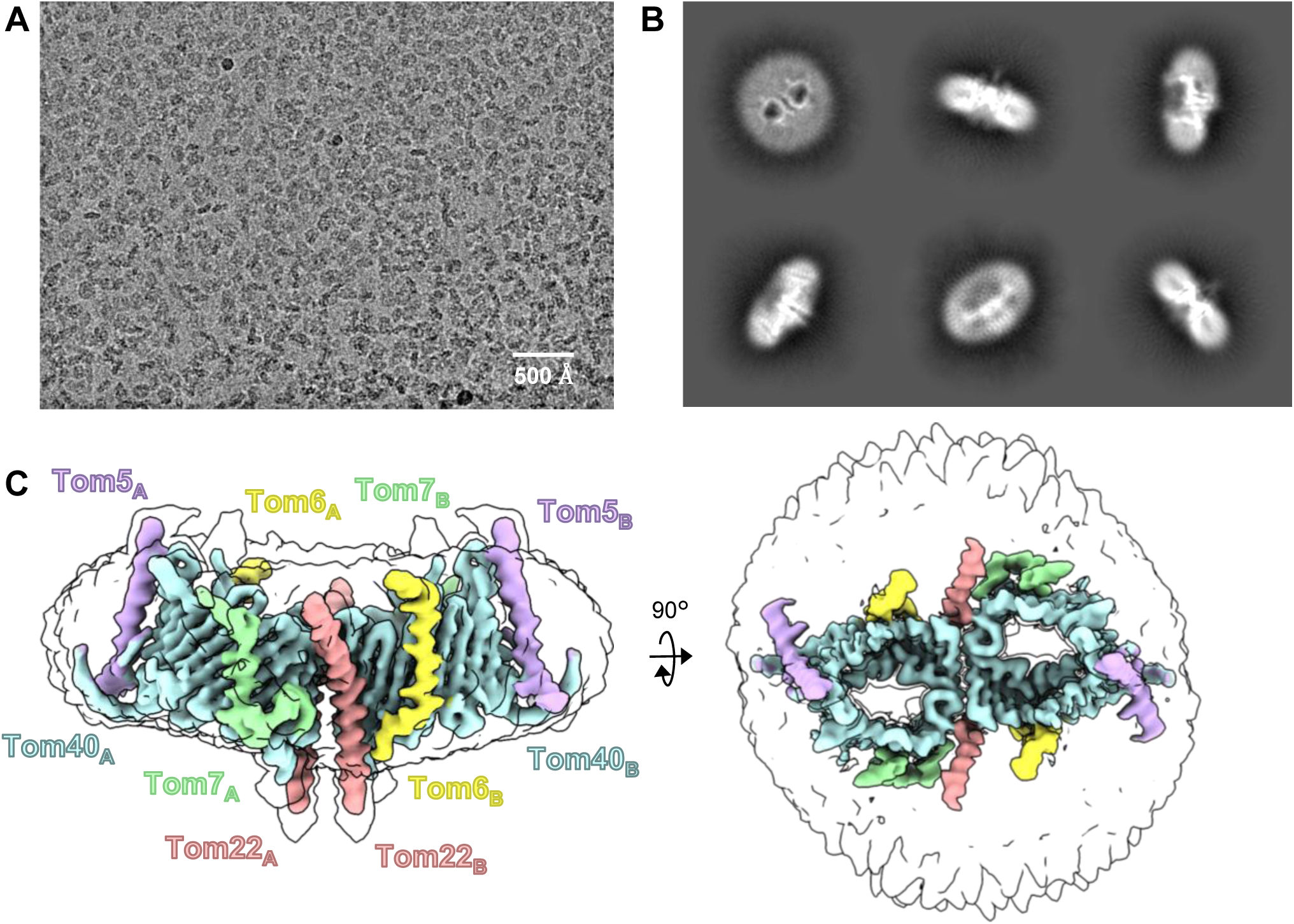
Representative HS-AFM images from independent observation of the TOM complex dissociation. The upper and lower images are independent. The trimers, dimers, monomers and substrates are indicated by blue, green, red, and yellow arrows, respectively. Imaging conditions were either 1.43 frames per second with a scan size of 200 × 200 nm^2^ and a pixel resolution of 320 × 80 on the top panel and 2.5 frames per second with a scan size of 120 × 84 nm^2^ and a pixel resolution of 400 × 70 on the bottom panel. The molecular images used in this figure were also cropped from the original data. The uncropped images were included in the supplementary figures.

**Figure S4.**
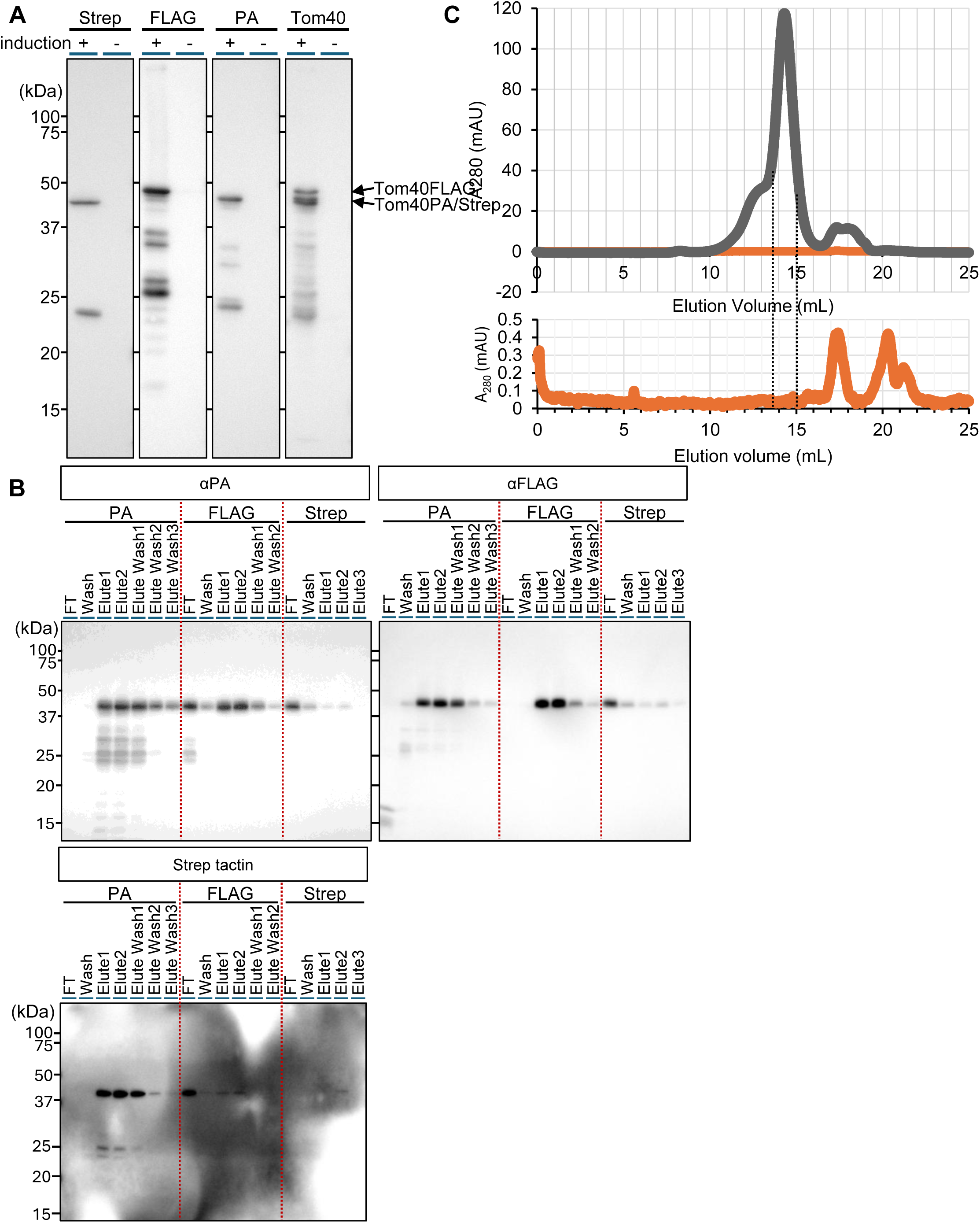
Purification of the triple-tagged TOM complex. (A) Evaluation of the expression levels of the Tom40 fused to the Strep II, 3×FLAG and PA -tags, respectively. (B) Western blot analysis for sample evaluation from independent purification of the triple-tagged TOM complex. (C) The gel filtration chromatograms of the TOM complex (upper, navy) and the triple-tagged TOM complex (lower, orange).

**Figure S5.**
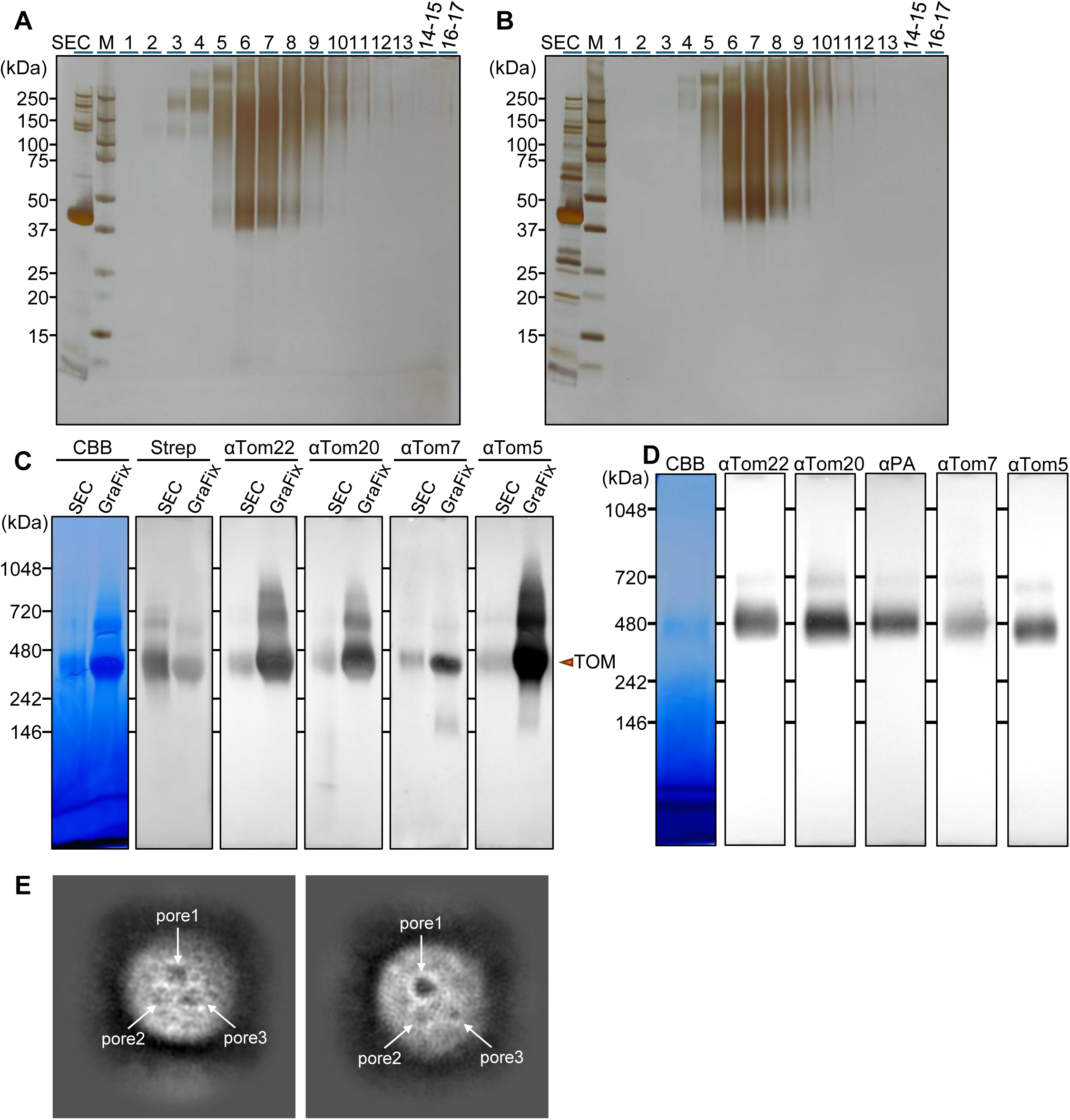
Sample evaluation of the crosslinked TOM complex by using GraFix method. SDS-PAGE analyses for the TOM complex (A) and the PA-TOM complex (B). BN-PAGE analyses for the TOM complex (C) and the PA-TOM complex (D). Representative 2D class average images of the same crosslinked TOM complex sample by GraFix method used for HS-AFM (E).

**Supplementary Table 1.**
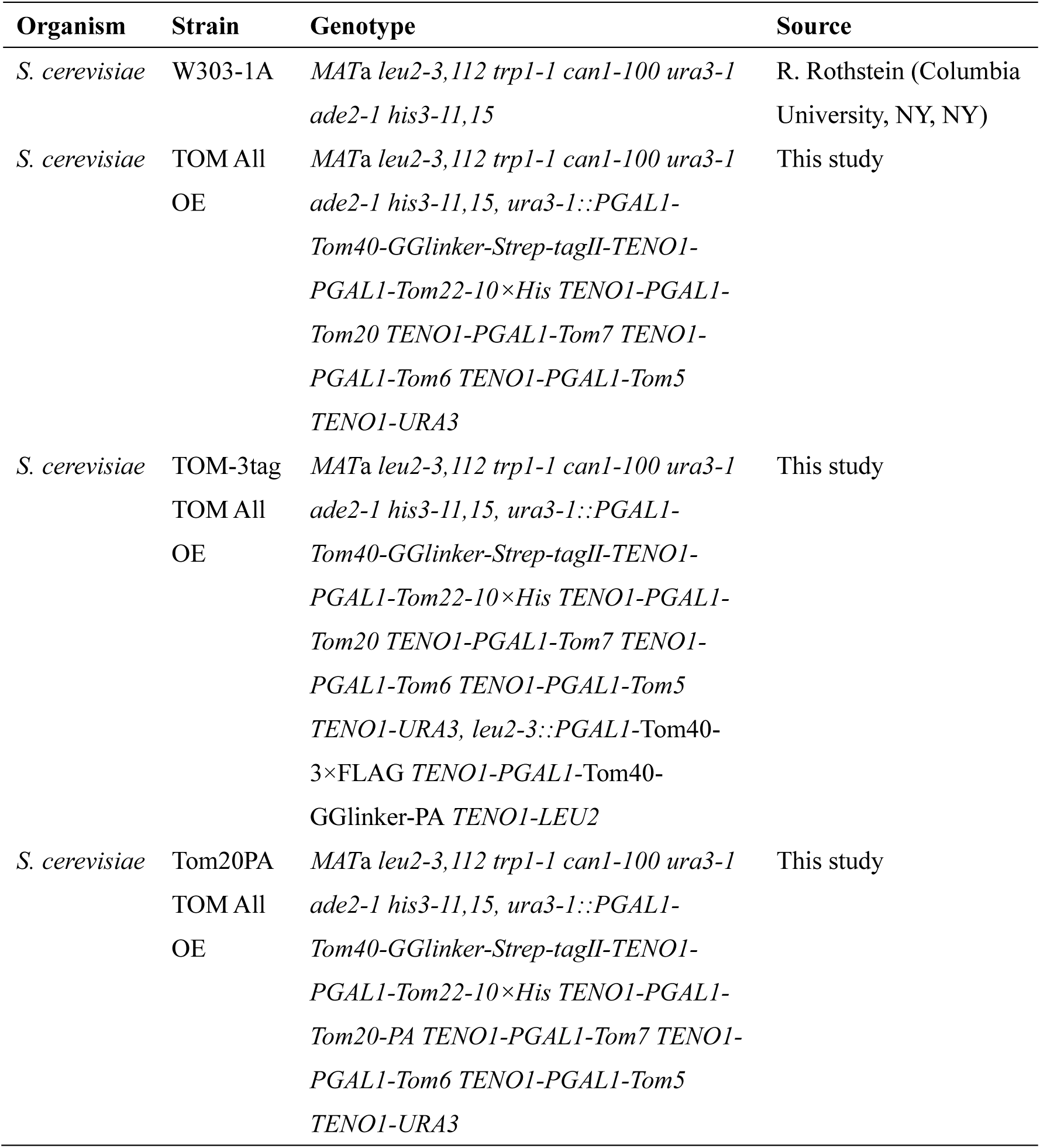
Yeast strains used in this study.

**Supplementary Table 2.**
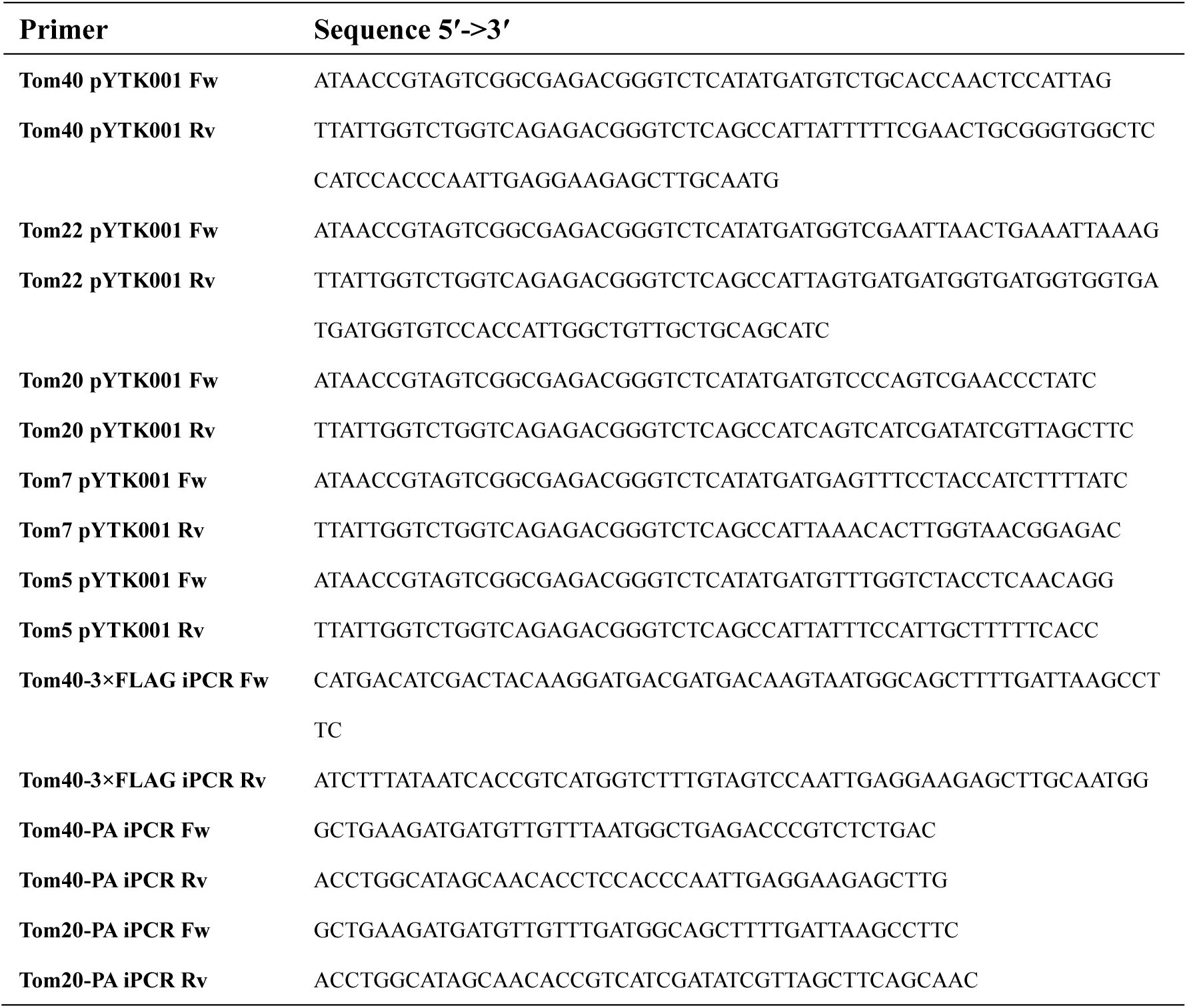
Primers used in this study.

**Supplementary Table 3.** Plasmids used in this study.

| Plasmid | Promoter | Expressed protein | Source |
| --- | --- | --- | --- |
| pYTK096/TOM complex | *GALI* | Tom40-GGlinker-Strep-tagII<br>Tom22-10×His<br>Tom20<br>Tom7<br>Tom6<br>Tom5 | This study |
| pYTK096(LEU2ver.)/TOM complex-Tom40-2tags | *GALI* | Tom40-3×FLAG<br>Tom40-GGlinker-PA | This study |
| pYTK096/TOM complex-Tom20PA | *GALI* | Tom40-GG-Strep-tagII<br>Tom22-10×His<br>Tom20-PA<br>Tom7<br>Tom6<br>Tom5 | This study |
| pET22b/pSu9-mDHFR | *T7* | pSu9-mouseDHFR | T. Endo<br>(Kyoto<br>Sangyo<br>University<br>, Kyoto,<br>Japan) |

**Supplementary Table 4.**
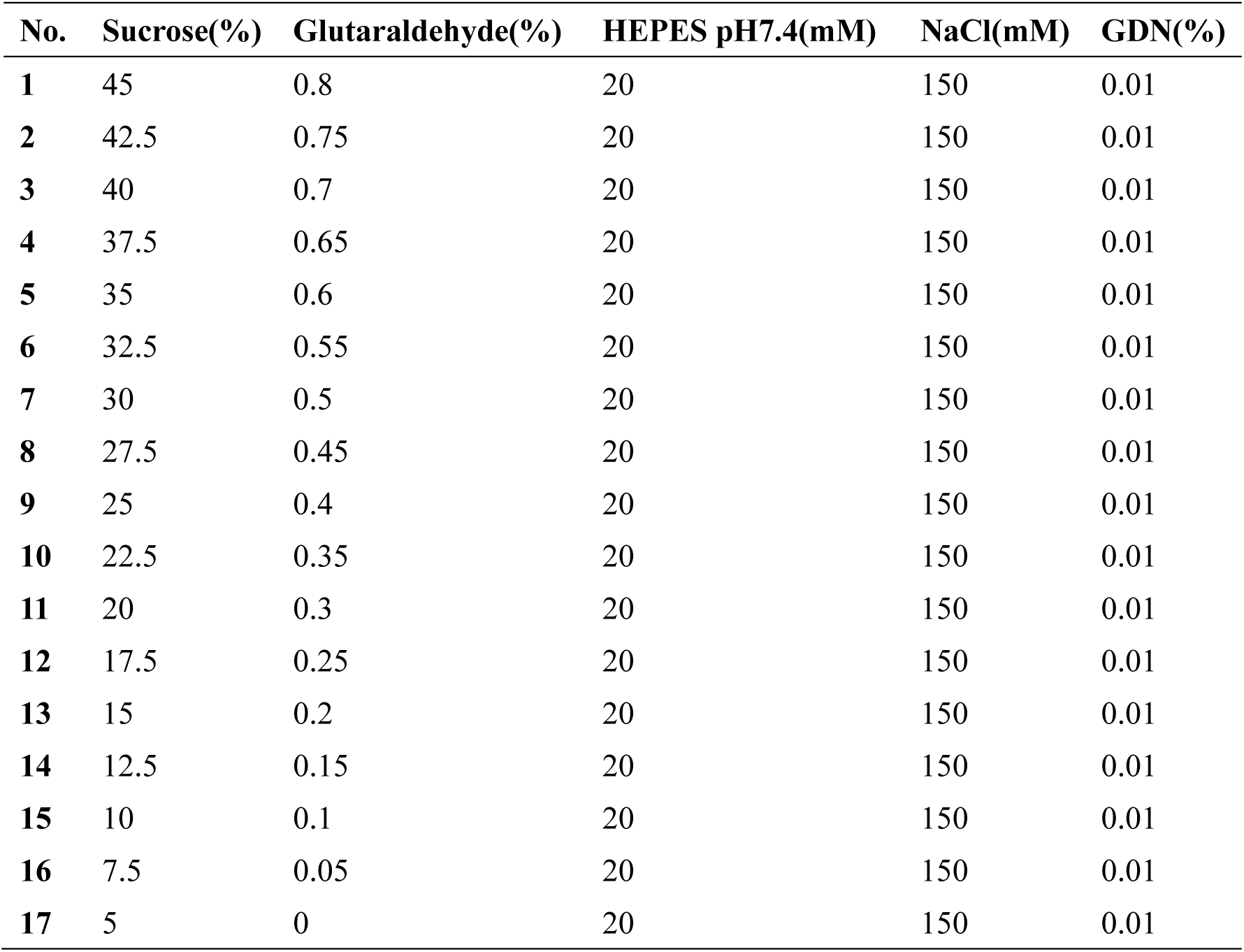
17-steps gradient buffer used in this study.

**Supplementary Table 5.** Data collection and processing for TOM complex.

| **Sample** | **TOM complex without cross-link** | **Cross-linked TOM complex** |
| --- | --- | --- |
| **Microscope** | Titan Krios G3i |  |
| **Detector** | Gatan K3 Camera |  |
| **Magnification** | 105,000 |  |
| **Voltage (kV)** | 300 |  |
| **Electron exposure (e<sup>-</sup>/Å<sup>2</sup>)** | 50 |  |
| **Defocus range (μm)** | −0.8 to −1.6 |  |
| **Pixel size (Å)** | 0.83 |  |
| **Symmetry imposed** | $C_1$ | |
| **Number of movies** | 8,397 | 4,189 |
| **Initial particle images (no.)** | 4,987,912 | - |
| **Final particle images (no.)** | 290,817 | - |
| **Map resolution (Å)** | 3.57 | - |
| **FSC threshold** | 0.143 |  |

